# Ammonium Transporters Govern Nutrient Sensing and Trap Induction in the Nematode-Trapping Fungi *Arthrobotrys oligospora*

**DOI:** 10.64898/2026.02.06.704490

**Authors:** Sheng-Chian Juan, Hillel T. Schwartz, Yen-Ping Hsueh

**Affiliations:** Institute of Molecular Biology, Academia Sinica, Taipei 115, Taiwan; Genome and Systems Biology Degree Program, Academia Sinica and National Taiwan University, Taipei 106, Taiwan; Department of Complex Biological Interactions, Max Planck Institute for Biology Tübingen, 72076 Tübingen, Germany

**Keywords:** nematode-trapping fungi, *Arthrobotrys oligospora*, ammonium transporter, nutrient sensing

## Abstract

To survive nutrient scarcity, organisms must detect environmental cues and acquire essential resources efficiently. Under nutrient-limiting conditions, nematode-trapping fungi (NTF) such as *Arthrobotrys oligospora* form specialized traps to capture nematodes; this transition can be suppressed by the ready availability of carbon and nitrogen. Ammonium transporters of the *MEP* family are key regulators of nitrogen sensing and uptake across all forms of life. We identified three *MEP* genes in *A. oligospora* based on protein sequence homology. Functional assays revealed that loss of the *MEP* genes alters the fungus’s ability to produce traps, without affecting the morphology of the traps or their ability to function in catching nematodes. Heterologous expression in *Saccharomyces cerevisiae* revealed functional conservation of Mep transporter subclasses. Together, the Mep transporters of *A. oligospora* contribute to ammonium-regulated trap induction.

## Introduction

Cells adapt to changing environments by sensing and responding to extracellular cues, often through membrane-associated nutrient sensors. Efficient assimilation of primary nutrients is essential for maintaining homeostasis and supporting growth, and is regulated at both transcriptional and post-translational levels. Nitrogen is a vital element, indispensable for the biosynthesis of nucleic acids and amino acids. To optimize nitrogen utilization, organisms employ nitrogen catabolite repression (NCR), a regulatory mechanism that promotes the use of preferred nitrogen sources such as ammonium and glutamine, while suppressing genes involved in the metabolism of less favorable sources such as urea and proline^1–3^. Ammonium, the most favored nitrogen source for many microorganisms, can be acquired either via nitrate reduction or direct uptake through ammonium transporters^4^.

Ammonium uptake is mediated by the Amt/Mep/Rh (Ammonium transporter/Methylammonium permease/Rhesus) protein family, which is conserved across all kingdoms of life^5^. To adapt to changing environments, many organisms express multiple ammonium transporters, which are typically classified into two groups according to their affinity and transport capacity for ammonium ^1^. One well-characterized example is the Mep transporter family in *Saccharomyces cerevisiae*, which includes three transporters: ScMep1 and ScMep3, comprising one group, exhibit high capacity but low affinity, whereas ScMep2 displays high affinity but low transport capacity. Increased expression of *ScMEP* genes is associated with the morphological transition from yeast to pseudohyphal growth under nitrogen-limiting conditions. In particular, *ScMEP2* expression is upregulated, and ammonium binding to ScMep2 activates the downstream MAPK pathway, promoting filamentous growth^6^. Deletion of *ScMEP2* abolishes this developmental response, indicating that ScMep2 functions as a transceptor—both transporting and sensing ammonium. Similar, filamentous growth in *Candida albicans* requires strong expression of the ammonium transporter gene *MEP2*, and filamentous growth of *Ustilago maydis* requires the ammonium transceptor Ump2, while overexpression of Ump2 causes filamentation even in the absence of ammonium deprivation^7,8^, underscoring the conserved role of ammonium transporters in morphogenesis and virulence.

Members of the Mep family of transporters share a conserved structure, typically containing 11–12 transmembrane domains and a conserved twin-His motif at the transport pore, with functional diversity stemming from key amino acid substitutions^9,10^. Substitution of the first of two histidines lining the pore of the *Arabidopsis* ammonium transporter AMT1-2 with glutamate decreased both the selectivity of the transporter and its affinity for ammonium, and increased transport rates^7,11,12^. In *S. cerevisiae*, both ScMep1 and ScMep3 carry this His-to-Glu substitution, which corresponds well with their increased ammonium transport capacity compared to ScMep2, which lacks this substitution^12,13^. A similar change in *Escherichia coli* AmtB converts the transporter into a channel-like conduit without structural disruption^10^, revealing the profound functional impact of a single residue change.

Nutrient availability is a key driver of lifestyle transitions in fungi. In the nematode-trapping fungus *Arthrobotrys oligospora*, nutrient depletion and prey signals trigger a transition from saprophytic to predatory growth. This developmental switch proceeds through distinct stages—prey attraction, sensing, trap development, capture, digestion, and conidiation—typically completed within 24 hours^14^. The fungus attracts nematodes via volatile compounds mimicking food and sex pheromones^15^, and detects nematode-derived ascarosides through G protein–coupled receptors, activating signaling pathways that initiate trap formation^16,17^. Supplementation with nitrogen or carbon suppresses this process, indicating that environmental nutrient levels directly regulate predatory behavior^16^. Time-course transcriptomic analyses have revealed stage-specific gene expression patterns, offering molecular insight into this transition^18^. Nonetheless, how *A. oligospora* senses, acquires, and integrates nutrient signals to regulate trap formation remains poorly understood.

In this study, we investigated the role of ammonium transporters in *Arthrobotrys oligospora*. Three putative ammonium transporter genes were identified and designated *MEP1*, *MEP2*, and *MEP3* based on protein sequence homology. Functional assays revealed that Mep1 and Mep3 contribute to the induction of trap formation, but not to trap morphogenesis or predation. Heterologous expression of *AoMEP* genes in yeast complemented mutants lacking the function of native ammonium transporters. Collectively, these findings demonstrate that ammonium transporters in *A. oligospora* play roles in regulating both hyphal growth and trap induction.

## Results

### *MEP1, MEP2*, and *MEP3* in *A. oligospora* are induced by nematodes and repressed by ammonium

Three homologs of the *Saccharomyces cerevisiae* ammonium transporter *MEP2* were identified in the *A. oligospora* genome and designated *MEP1*, *MEP2*, and *MEP3*, based on protein sequence similarity (Figure 1A). Phylogenetic analysis revealed that AoMep1 clusters with low-affinity, high-capacity transporters including ScMep1 and ScMep3, whereas AoMep2 and AoMep3 group with high-affinity transporters including ScMep2.

**Figure 1.**
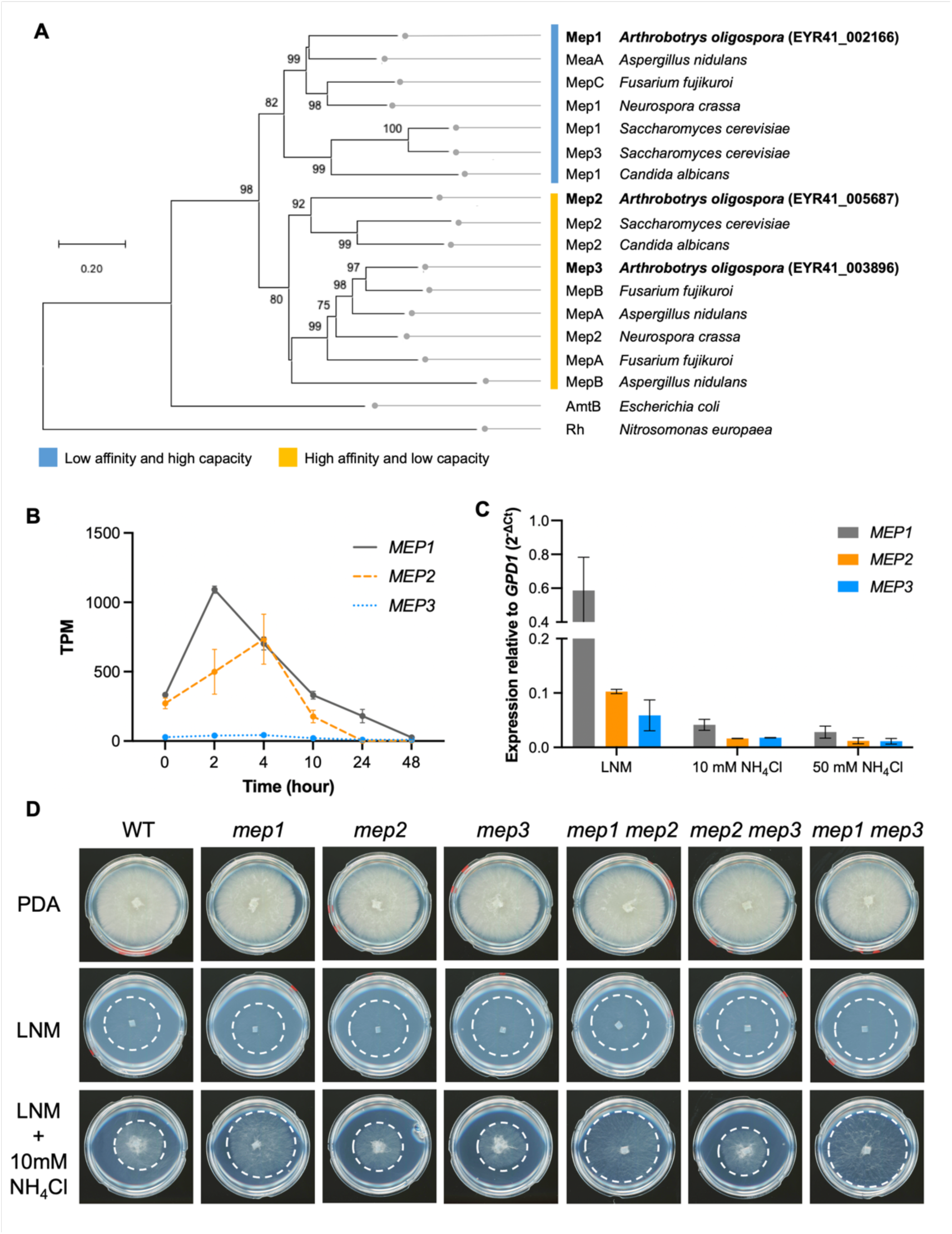
Three Mep ammonium transporters were characterized in *A. oligospora*. (A) Phylogenetic analysis of fungal ammonium transporters was performed in MEGA11^32^ using the Neighbor-Joining method with 3000 bootstrap replicates. The tree is drawn to scale, with branch lengths reflecting evolutionary distances calculated using the JTT matrix-based method.^33^. The blue bar indicates low-affinity, high-capacity transporters, while the yellow bar indicates high-affinity, low-capacity transporters, as determined by our analysis and prior studies. Accession numbers for reference transporters are listed in Supplementary Table 1. (B) *MEP* gene expressions in the time-course RNA-Seq from 0 to 48 hours post exposure to nematodes. TPM, transcripts per million reads. (C) *MEP* gene expression in wildtype examined by quantitative PCR under LNM and LNM with 10 mM or 50mM ammonium supplementation. (D) Hyphal growth on different media. Growth of all mutants was comparable to that of the wild type on either nutrient-rich medium (PDA) or nutrient-depleted (LNM) medium, while mutants lacking *MEP1* showed a partial ability to tolerate an elevated level of ammonium that inhibited the growth of the wild type. WT, wild type.

Most of the described motifs, key residues, and predicted transmembrane domains of previously studied ammonium transporters were found to be identical or in some cases highly conserved in the AoMep proteins (Supplementary Figure 1). Structural predictions indicate that the core structures of AoMep transporters are well aligned with the corresponding Mep transporters in *S. cerevisiae*. (Supplementary Figure 2). Notably, AoMep1 carries a Glu-to-His substitution at the first position of the conserved twin-His motif within transmembrane helix V, whereas AoMep2 and AoMep3 retain the canonical histidine at this site. Similar substitutions have been reported in other organisms that encode multiple ammonium transporters and are associated with altered transport properties^9,13,19–21^, suggesting potential functional diversification among Mep family members in *A. oligospora*.

To determine whether *MEP* gene expression in *A. oligospora* is modulated by nematode exposure and nitrogen availability, we analyzed transcript levels during predation and under varying ammonium conditions. In a time-course transcriptomic analysis, *MEP1* was rapidly induced upon exposure to *C. elegans*, peaking at 2 hours before gradually declining. *MEP2* showed delayed induction, with peak expression at 4 hours, while *MEP3* remained consistently low throughout the predation process (Figure 1B). These distinct temporal patterns suggest differential regulation of ammonium transporters in response to nematode cues. We next assessed *MEP* gene expression under varying ammonium concentrations using quantitative PCR. In nitrogen-free medium, *MEP1* showed the highest expression, followed by *MEP2* and *MEP3*. Supplementation with 10 or 50 mM ammonium suppressed expression of all three genes (Figure 1C). Collectively, these data show that *MEP* genes are differentially induced during predation and broadly repressed by elevated ammonium.

### Mep1 and Mep3 regulate trap formation in *A. oligospora*

To investigate the role of Mep transporters in fungal predation, we generated single and double gene deletion mutants by homologous recombination (Supplementary Figure 3). A triple *MEP* deletion mutant could not be constructed due to limitations in available selectable drug resistance markers. Under both nutrient-rich (PDA) and nutrient-limited (LNM) conditions, *mep* mutants and double mutants displayed colony expansion comparable to the wild-type strain (Figure 1D).

We then quantified trap formation in wild-type and *mep* mutant strains in response to their nematode prey. Under nutrient-limited conditions, the wild type produced over 400 traps per cm². Both *mep1* and *mep3* single mutants showed a significant reduction in trap formation, generating approximately half the number of traps, while the *mep2* mutant formed trap numbers comparable to the wild type. All *mep* double mutants were mutant for *mep1* or for *mep3*, or both, and showed trap formation levels comparable to *mep1* and *mep3* single mutants (Figure 2A, 2B). We further evaluated trap function using a nematode survival assay under continuous nematode exposure. After 12 hours, ∼25% of nematodes survived in the presence of wildtype *A. oligospora* or mutants only lacking *MEP2*, whereas survival rates exceeded 50% in the presence of mutants lacking *MEP1* or *MEP3* (Figure 2C). These results indicate that Mep1 and Mep3 are important for trap induction, such that the loss of either Mep1 or Mep3 reduces the fungus’s ability to trap and kill nematodes, whereas Mep2 appears to be dispensable for these processes.

**Figure 2.**
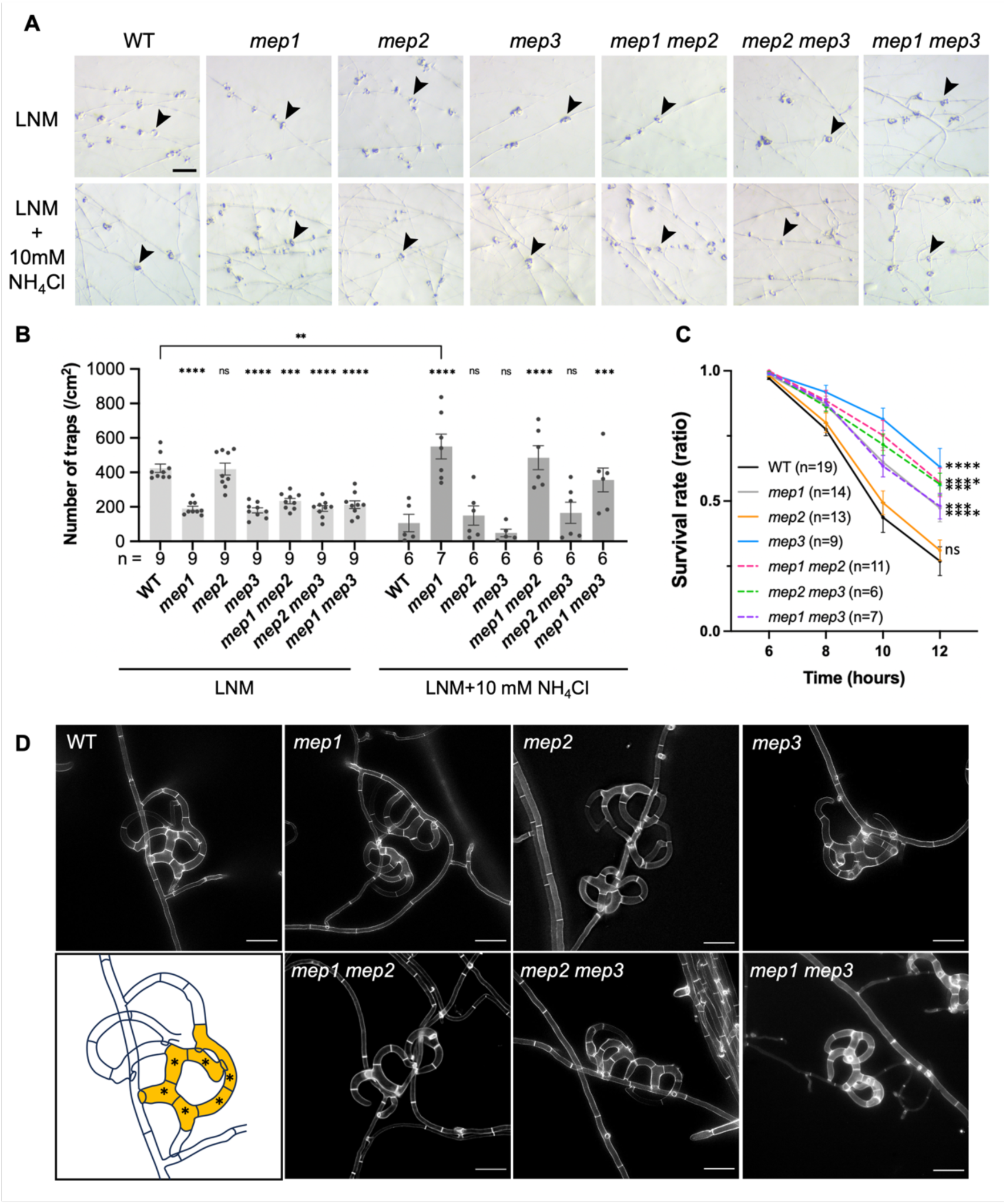
Phenotypic assays of *mep* mutants. (A) Representative brightfield images of the traps induced by nematodes in the WT and *mep* mutants on LNM or 10 mM ammonium chloride supplementation. Arrowheads indicate representative trap structures. WT, wild type. Scale bar, 2 mm.(B) Quantification of trap formation in *(A)*. Data are presented as mean ± SEM. Comparisons within groups were made with the wildtype and were analyzed by two-way ANOVA followed by Dunnett’s multiple comparisons test (\*\*\**P* < 0.0002, \*\*\*\**P* < 0.0001). Comparisons between groups were analyzed by two-way ANOVA followed by Sidak’s multiple comparisons test (\*\**P* < 0.01). (C) Survival rate assay of nematodes in response to wild-type and mutant strains. Nematode survival rates were higher upon exposure to most *mep* mutants than to the wild type, except for *mep2*. Significance was analyzed at the last timepoint using two-way ANOVA followed by Dunnett’s multiple comparisons test (\*\*\**P*<0.0002, \*\*\*\**P*<0.0001). (D) Examples of traps produced by each strain, with fungal cell walls stained by SR2200. The drawing takes the wild-type as an example and illustrates the method used to quantify the number of cells in the trap. A single loop is indicated in yellow, and the asterisks represent the counted cells in that loop. The quantification data is shown in Figure S3C. Scale bars, 30 μm.

### *mep1* mutants gain tolerance to ammonium

Ammonium is known to suppress trap formation in *A. oligospora*^16^. When 10 mM ammonium was added to LNM, the wild-type strain formed smaller colonies with densely packed hyphae compared to LNM condition. In contrast, *mep1*-deficient mutants produced larger colonies with more loosely arranged hyphae compared to the wild-type (Figure 1D). Adding ammonium to low-nitrogen media significantly reduced the number of traps that the wild type produced in response to nematodes, while *mep1* mutants produced as many or more traps under ammonium-rich conditions as the wild type did on low-nitrogen media (Figure 2A, B). On 50 mM ammonium media, both colony growth and trap formation were further suppressed in the wild-type compared to on LNM or on LNM supplemented with 10 mM ammonium, while *mep1* mutants continued to show better growth (Supplementary Figure 4A, B). Taken together, these results indicate that deletion of *mep1* confers partial tolerance to elevated levels of ammonium, highlighting its role in mediating the fungal response to ammonium availability.

### *mep* mutants produce fully functional traps with normal morphogenesis

To determine whether trap architecture differs between the wild-type and *mep* mutants, we examined trap morphology in detail. Traps formed by the *mep* mutants exhibited the normal multi-loop, net-like structures observed in the wild type. We also quantified the number of cells per trap loop as an indicator of trap morphogenesis. In the wild type, most traps contained four to seven cells per loop (mean ± SEM: 5.28 ± 0.15; n = 25). The number of cells per loop did not differ significantly from the wild type in any *mep* single or double mutant (Figure 2D; Supplementary Figure 4C), indicating that Mep transporters are not major regulators of trap morphogenesis.

We next evaluated trap function using a nematode capture assay. All strains captured nearly all the nematodes within 10 minutes (Supplementary Figure 4D), suggesting no defect in trap adhesiveness. Combined with earlier results, these findings indicate that Mep transporters influence trap induction but are dispensable for trap structure and function.

### *MEP* genes exhibit condition-dependent cross-regulation in *A. oligospora*

Functional redundancy among ammonium transporters has been reported in various fungi^11,22–24^. Consistent with this, *A. oligospora* double *mep* mutants did not display stronger phenotypes than the corresponding single mutants (Figure 2B), suggesting potential cross-regulation among *MEP* genes. To test this possibility, we quantified *MEP* transcript levels in the *mep* mutants by qPCR under different nitrogen conditions and compared them with wild type. In LNM, *mep1* and *mep2* mutants showed little change in expression of the remaining *MEP* genes, whereas the remaining *MEP* genes were modestly downregulated in the *mep3*, *mep1 mep2*, and *mep2 mep3* mutants (Supplementary Figure 5A). In contrast, supplementation with 10 mM NH₄Cl triggered a clear compensatory response: transcript levels of the remaining *MEP* genes increased to wild-type levels or higher, with particularly strong induction in backgrounds lacking *MEP1* (Supplementary Figure 5B), indicating condition-dependent transcriptional compensation among ammonium transporters. The marked upregulation of *MEP3* in *mep1* and *mep1 mep2* strains suggests that *MEP3* serves as a key alternative ammonium transporter when primary transport capacity is compromised.

### AoMep transporters harbor conserved functions as ScMep transporters

To assess the functional conservation of Mep transporters, we heterologously expressed *AoMEP* genes in *S. cerevisiae mep* mutants. *AoMEP* cDNAs were cloned into constructs containing the constitutive *PGK1*promoter in the high-copy plasmid pRS42N to ensure robust expression (Supplementary Figure 6).

To examine whether AoMep transporters can function in ammonium sensing, we expressed each *AoMEP* gene in a *S. cerevisiae mep2* mutant (MLY108a/α), a commonly used system to evaluate ammonium-sensing ability based on rescue of pseudohyphal growth under low-ammonium conditions (Figure 3A)^22^. Expression of *AoMEP2* or *AoMEP3* fully restored pseudohyphal growth in the *Scmep2* mutant, whereas *AoMEP1* supported only limited filamentation. Given that AoMep2 and AoMep3 are phylogenetically clustered with ScMep2, this result is consistent with its predicted functional class.

**Figure 3.**
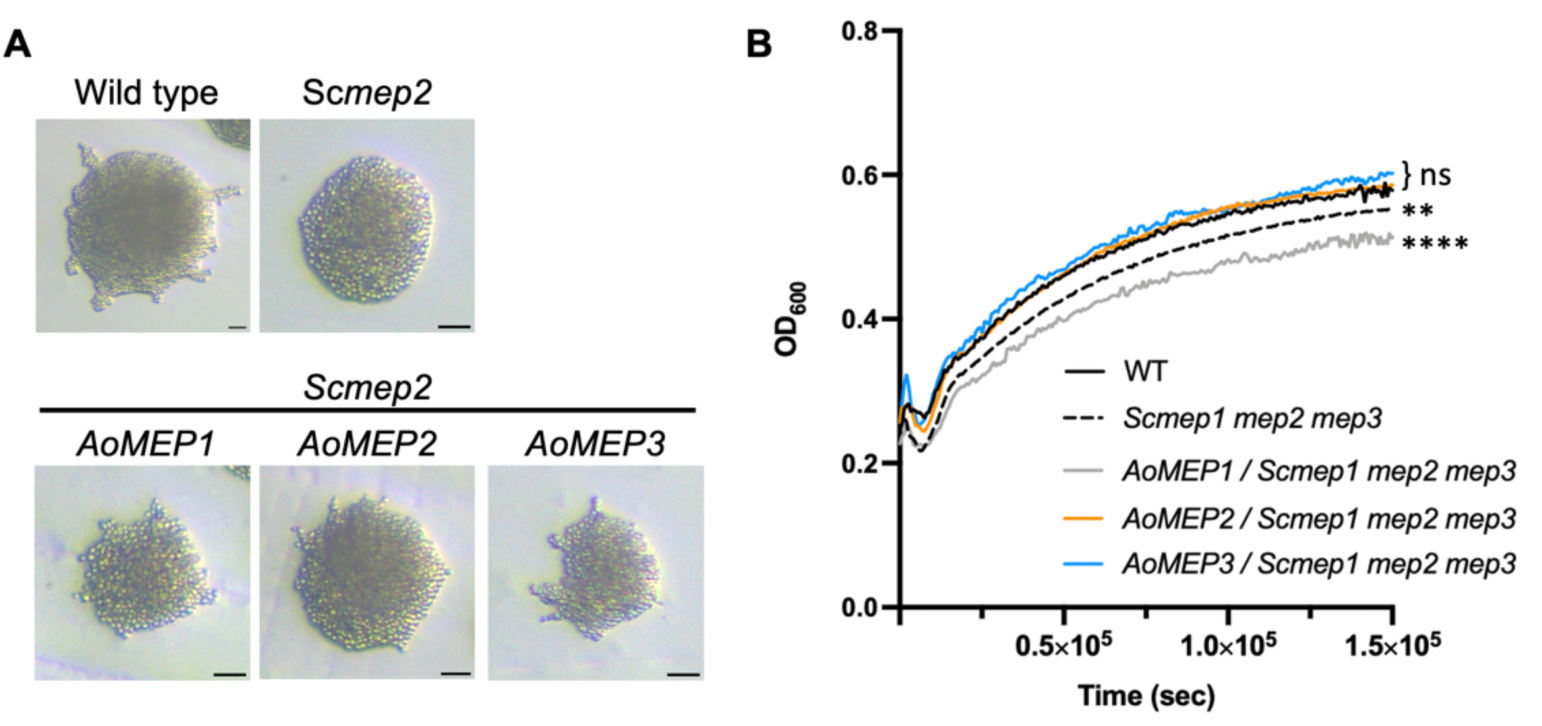
Heterologous gene expression of *AoMEP* genes in *S. cerevisiae* and functional assays. (A) Pseudohyphal growth assay of wildtype *S. cerevisiae*, the *Scmep2* mutant, and the *Scmep2* mutant expressing the indicated *AoMEP* genes grown on SLADG agar medium supplemented with 25 μM (NH_4_)_2_SO_4_ at 30°C for 6 days. *AoMEP1* partially complemented pseudohyphal growth, whereas *AoMEP2* and *AoMEP3* fully complemented pseudohyphal growth. Scale bar, 0.1 mm. (B) *S. cerevisiae Scmep1 mep2 mep3* (MLY131a/*⍺*) triple mutant growth defect complementation assay. *AoMEP* genes were transformed individually into the *Scmep1 mep2 mep3* mutant and grown in SLADG with 25 μM (NH_4_)_2_SO_4_ at 28°C for 2 days. *AoMEP2-* and *AoMEP3-*transformed triple mutants were rescued for its slow growth phenotype and were not statistically different from the wild type, while the *AoMEP1-*transformed *Scmep1 mep2 mep3* triple mutant grew even more solely than the untransformed mutant. Statistical significance was determined by one-way ANOVA followed by Dunnett’s multiple comparisons test (\*\**P*<0.002, \*\*\*\**P*<0.0001).

We next examined whether *AoMEPs* could rescue the growth defect of the *Scmep1Δmep2Δmep3Δ* triple mutant (MLY131a/α) under low nitrogen conditions^22^. In SLAD medium containing 25 μM ammonium sulfate, *Scmep1Δmep2Δmep3Δ* triple mutant strains expressing *AoMEP2* or *AoMEP3*-expressing strains displayed growth curves comparable to the wild type, whereas *AoMEP1* expression failed to restore growth to the *Scmep1Δmep2Δmep3Δ* triple mutant and even enhanced its growth defect (Figure 3B). Together, these findings indicate that AoMep2 and AoMep3 can functionally substitute for *S. cerevisiae* Mep transporters in both ammonium sensing and uptake, supporting evolutionary conservation of Mep transporter function across fungal species.

## Discussion

In this study, we characterized three ammonium transporter genes in the nematode-trapping fungus *A. oligospora* — *AoMEP1*, *AoMEP2*, and *AoMEP3* — and found that Mep-mediated ammonium transport contributes to the regulation of predatory development. All three AoMep proteins retain conserved features of the Amt/Mep family, including predicted transmembrane architecture and signature residues required for ammonium transport. Phylogenetic placement identified indications of functional diversification within the AoMep family: AoMep1 clusters with low-affinity/high-capacity transporters including ScMep1 and ScMep3, while AoMep2 and AoMep3 are grouped with high-affinity/low-capacity transporters such as ScMep2. Consistent with this division, heterologous expression in *S. cerevisiae* supported functional conservation of these transporter subclasses: expression of *AoMEP2* or *AoMEP3* in a *S. cerevisiae* strain lacking all three Mep ammonium transporters complemented the mutant strain’s defects in ammonium-dependent pseudohyphal growth under limiting ammonium and growth on low-nitrogen medium, while expression of *AoMEP1* provided only partial complementation of pseudohyphal growth and did not restore growth on low-nitrogen medium.

One notable structural feature distinguishing AoMep1 from AoMep2/AoMep3 is the His-to-Glu substitution at the first position of the conserved twin-His motif (Supplementary Figure 1). Similar substitutions have been shown in other systems to alter transport properties and can shift a transporter toward channel-like behavior or increased transport capacity^10,13^. In *A. oligospora*, deletion of *AoMEP1* conferred increased tolerance of elevated ammonium, improving colony expansion and reducing suppression of trap formation. These phenotypes were not observed in *mep2* or *mep3* mutants, suggesting that AoMep1 mediates the cellular response to high external ammonium, likely by transporting ammonium into the cell.

Our results further indicate that AoMep transporters primarily influence trap induction, rather than trap morphogenesis or trap function. Mutants lacking *MEP1* or *MEP3* formed significantly fewer traps and showed reduced killing efficiency in survival assays, yet the traps that formed had normal architecture and retained the ability to rapidly and efficiently capture their nematode prey, indicating that after Mep transporters influence the decision to produce traps, they are not required for the development of functional traps. This suggests that AoMep-dependent ammonium uptake and sensing acts upstream of the developmental decision to enter predatory growth, rather than controlling the morphogenetic machinery itself.

Redundancy and compensatory regulation among ammonium transporter paralogs have been described in many organisms. Consistent with this, *A. oligospora* double *mep* mutants did not show more severe defects than the corresponding single mutants, suggesting that loss of one transporter can be buffered by others. Our qPCR results support this idea by revealing condition-dependent cross-regulation among *MEP* genes. Under nitrogen limitation, expression of the remaining *MEP* genes changed only modestly. In contrast, ammonium supplementation triggered clear compensatory induction of the remaining *MEP* genes, with particularly strong upregulation in strains lacking *MEP1*. Together, these data indicate that *MEP* paralogs can be transcriptionally rewired in response to transporter loss—especially in ammonium-rich conditions—thereby helping maintain overall ammonium uptake capacity and explaining why double mutants do not exhibit markedly stronger phenotypes.

Together, our results support a model in which the nematophagous fungus *A. oligospora* uses Mep ammonium transporters to integrate environmental nitrogen availability with prey-derived signals to regulate the initiation of predatory development (Supplementary Figure 7). Under low-ammonium conditions, the high-affinity transporters AoMep2 and AoMep3 uptake of scarce ammonium; however, overall nitrogen limitation persists, a condition that allows the fungus to respond to nematode-derived cues by initiating robust trap production. By contrast, under high-ammonium conditions the low-affinity, high-capacity transporter AoMep1 mediates ammonium uptake, providing sufficient intracellular ammonium levels to activate downstream nitrogen-sufficiency signaling, thereby attenuating starvation-associated signaling and suppressing trap formation even in the presence of prey cues. In this framework, AoMep transporters act as part of a nutrient-sensing module that responds to environmental nitrogen conditions to govern the developmental switch between saprophytic growth and predatory behavior.

As nematode-trapping fungi have been considered as potential biocontrol agents against plant-parasitic nematodes^25^, which cause substantial agricultural losses worldwide^26,27^, one limitation is that nutrient-rich agricultural environments can suppress the predatory transition. This study provides insight into how *A. oligospora* senses and responds to environmental nitrogen availability via Mep ammonium transporters, thereby linking ammonium status to trap induction. These findings suggest practical ways to improve the reliability of NTF-based biocontrol in agricultural settings, for example by adjusting fertilizer practices to minimize ammonium-mediated suppression of trap formation, or by selecting or developing fungal strains that are less sensitive to high nitrogen conditions, thereby improving control of nematodes and reducing reliance on chemical nematicides.

## Material and methods

### Strains and media

The *A. oligospora* strains used in this study are listed in Table 1. All mutations were introduced into the *ku70* background to improve gene knockout efficiency via homologous recombination^28^. The cultivation of *A. oligospora* was carried out on potato dextrose agar (PDA), potato dextrose broth (PDB), and low-nutrient medium (LNM, 2% agar, 1.66 mM MgSO_4_, 5.4 µM ZnSO_4_, 2.6 µM MnSO_4_, 18.5 µM FeCl_3_, 13.4 mM KCl, 0.34 µM biotin, and 0.75 µM thiamin) at 25°C.

**Table 1.** *Arthrobotrys oligospora* strains used in this study.

| Strain | Strain no. | Genotype | Reference |
| --- | --- | --- | --- |
| **Wild type** | TWF154 | - | (Yang, *et al.*, 2020) |
| ***ku70\**** | TWF1697 | $\Delta ku70::hph$ | (Kuo, Chen, & Hsueh, 2020) |
| ***mep1*** | TWF3420 | $\Delta ku70::hph \Delta mep1::nat1$ | This study |
| ***mep2*** | TWF3653 | $\Delta ku70::hph \Delta mep2::neo$ | This study |
| ***mep3*** | TWF3717 | $\Delta ku70::hph \Delta mep3::nat1$ | This study |
| ***mep1 mep2*** | TWF3655 | $\Delta ku70::hph \Delta mep1::nat1 \Delta mep2::neo$ | This study |
| ***mep2 mep3*** | TWF3703 | $\Delta ku70::hph \Delta mep3::nat1 \Delta mep2::neo$ | This study |
| ***mep1 mep3*** | TWF3932 | $\Delta ku70::hph \Delta mep1::nat1 \Delta mep3::neo$ | This study |

\*The *ku70* strain is used as the wild-type experimental control for all assays throughout this study.

For transporter functional assays using *Saccharomyces cerevisiae* strains, refer to Supplementary Table 3. Yeast cultures were maintained at 25°C either in liquid or solid yeast extract-peptone-dextrose (YPD) medium, consisting of 1% (w/v) yeast extract, 2% (w/v) peptone, and 2% (w/v) dextrose. Pseudohyphal growth complementation assays were performed on synthetic low ammonium dextrose galactose agar (SLADG), which contained 0.17% (w/v) yeast nitrogen base without amino acids or ammonium sulfate, 0.2% (w/v) dextrose, 2% (w/v) galactose, 2% (w/v) agar, and 50 μM ammonium sulfate, supplemented with 20 μg/mL uracil and 120 μg/mL leucine. Growth defect complementation assays were carried out using SLADG and synthetic dextrose medium (SD), composed of 0.67% (w/v) yeast nitrogen base without amino acids, 2% (w/v) dextrose, and 2% (w/v) agar, also supplemented with 20 μg/mL uracil and 120 μg/mL leucine.

The *Caenorhabditis elegans* strain used in this study was the wild-type strain N2 (Bristol), which was maintained on nematode growth medium (NGM) plates and fed with *Escherichia coli* strain OP50^29^.

### Bioinformatic analyses

The amino acid sequence of Mep2 from *S. cerevisiae* was used as a query to identify homologous proteins in *A. oligospora* using local BLAST searches implemented in Blast2GO 5 PRO^30^. Protein sequence alignments between AoMep and ScMep transporters were generated using MAFFT (v7)^31^. Phylogenetic analysis of ammonium transporters from selected fungal species (Supplementary Table 1) was performed in MEGA11^32^ using the Neighbor-Joining method with 3000 bootstrap replicates. Evolutionary distances were calculated using the JTT matrix-based model.^33^. Predicted transmembrane helices of Mep proteins were identified using TMHMM v2.0^34^.

### Protein structure prediction and alignment

Structural prediction of AoMep proteins was carried out using AlphaFold^35^. Structural similarity between AoMep transporters and *S. cerevisiae* Mep transporters was evaluated using Foldseek-based structural alignment^36^. Structural alignments were visualized and rendered using PyMOL^37^.

### Generation of the DNA cassette for gene deletion

The primers used to generate gene deletion and complementation constructs are listed in Supplementary Table 2. To perform gene deletion via homologous recombination, the 5’ and 3’ homologous flanking regions of each target gene were amplified using PCR from the total genomic DNA of the *A. oligospora* TWF154 strain using overlapping primers (as shown in Supplementary Figure 3). The selection marker-resistant genes *NAT1* and *GentR* were amplified from the plasmids pRS41N^38^ and pUMa1057^39,40^, respectively. These flanking regions were fused with the drug-resistant gene placed in the middle, using overlap polymerase chain reaction (PCR). The resulting construct was further amplified through nested PCR to ensure the successful assembly of the gene deletion cassette.

### Generation of target gene deletion mutant strains

The protoplast preparation was based on the established protocol^41^. In brief, fungal hyphae of the *A. oligospora ku70* strain (TWF1697) or of mutants of *A. oligospora* were cultured in potato dextrose broth (PDB) for 1 day and harvested by centrifugation at 5000 rpm. The hyphae were then rinsed with MN buffer (0.3 M MgSO₄, 0.3 M NaCl) and enzymatically digested into protoplasts using 50 mg/mL VinoTastePro (Novonesis) in MN buffer, incubating at 25°C for 15-17 hours. The resulting protoplasts were filtered through sterile Miracloth (Millipore) and centrifuged at 1500 rcf for 5 minutes. Afterward, the protoplasts were washed with STC buffer(1.2 M sorbitol, 50 mM CaCl2, 10 mM Tris– HCl pH 7.5) and suspended in fresh STC buffer, with their concentration determined using a hemocytometer (Mariendeld Superior).

For each transformation, 3 × 10⁵ protoplasts were mixed with 2.5 μg of the DNA construct in a 50 mL centrifuge tube and incubated on ice for 30 minutes. After incubation, five volumes of PTC solution (40% w/v PEG4000, 10 mM Tris-HCl [pH 7.5], 50 mM CaCl₂) were gently added, and the mixture was incubated at room temperature for 20 minutes. To facilitate protoplast regeneration, regeneration medium (0.3% w/v acid-hydrolyzed casein, 0.3% w/v yeast extract, 20.5% w/v sucrose, 1% w/v agar) was added to a final volume of 50 mL. The medium was supplemented with either 180 μg/mL of nourseothricin (clonNAT) or 250 μg/mL of geneticin (G418), depending on the selection marker present in the DNA construct. The mixture was gently combined and evenly spread across three empty 9 cm Petri dishes, which were then incubated at 25°C for one week to allow for colony formation and selection.

Transformants that emerged on the regeneration plates were subsequently transferred to PDA Petri plates containing the appropriate selection drug for further screening. These potential transformants were then cultured on PDA at 25°C to ensure stability and for use in subsequent experimental procedures.

### Gene deletion validation

To verify gene deletion, the DNA of potential mutants was confirmed via PCR using the primers listed in Supplementary Table 2. Internal primers were designed to bind within the target gene, while external primers targeted the 5’ flanking region and the selection drug resistance gene. Mutants lacking the target gene showed no internal band, and the expected sizes of external bands were used to confirm successful deletion before proceeding with subsequent assays.

For pure genomic DNA isolation from *A. oligospora*, CTAB-PVP-based DNA extraction method followed by chloroform purification was employed as previously described^42^. After extraction, genomic DNA was further purified using the Genomic DNA Clean & Concentrator-10 kit (Zymo Research). Genomic DNA was first checked by PCR before Southern blotting.

Digoxigenin (DIG)-based Southern blotting was performed to check for possible ectopic insertions of the target gene. In brief, total genomic DNA from wildtype and mutant strains was purified and digested using the restriction enzymes specified in Supplementary Figure 3. The digested DNA samples were separated on a 0.9% agarose gel and transferred onto a nylon membrane overnight for subsequent analysis. DIG-labeled probes were generated using PCR with a PCR DIG Probe Synthesis Kit (Roche). The nylon membrane was hybridized overnight with a single-stranded DIG-labeled probe. Following the hybridization, the membrane underwent a series of washing steps to remove non-specific binding. The membrane was then incubated with alkaline phosphatase-coupled anti-DIG antibody (Anti-Digoxigenin-AP, Fab fragments, Roche). Detection of DIG-labeled fragments was achieved using CDP-Star solution (Roche), which enabled visualization of the hybridized probes.

### Trap number quantification

To assess fungal sensitivity to nematodes, all strains were cultured on 3 cm Petri plates containing low-nutrient medium (LNM) or LNM supplemented with either 10 mM or 50 mM NH₄Cl at 25°C for 2 days. After this incubation period, 60 *C. elegans* N2 larvae at the fourth larval (L4) stage were added to each plate to induce trap formation. Following a 6-hour exposure, the nematodes were removed by washing the plates with distilled water (ddH₂O). The plates were then incubated at 25°C without Parafilm to continue the experiment. 24 hours after nematode addition, three random positions on each plate were photographed using a ZEISS Stemi 305 Stereo Microscope at 40× magnification. The total number of traps observed in the three photos was normalized to represent the trap count per cm², which was used as the trap number for each plate. This quantification allowed for a comparison of trap formation across different strains and ammonium conditions.

### Caenorhabditis elegans survival rate assay

All *A. oligospora* strains were cultured on 3 cm LNM Petri plates and incubated at 25°C for 2 days. Then, 80 young-adult stage N2 nematodes were added onto the plate. After 6 hours, the numbers of crawling nematodes were counted every 2 hours for a total of 12 hours. Survival rate was calculated by dividing the number of living (motile) nematodes by 80.

### Capture rate assay

To test if the traps generated by fungi were functional, all strains were cultured on 5 cm LNM plates at 25°C for 3 days. Then, 200 young-adult stage N2 nematodes were added to induce trap formation overnight. The next day, 30 N2 adults were placed onto the plate for 10 minutes, before adding 1 mL ddH_2_O to quantify the number of un-trapped nematodes. Capture rate was calculated by dividing the number of living (motile) nematodes by 30.

### Imaging of trap structures and cell number quantification

Fungi were grown on LNM containing 0.1% SCRI Renaissance Stain 2200 (SR2200, Renaissance Chemicals Ltd) for cell wall-specific staining. After nematode induction, images of traps were taken using a ZEISS AxioObserver Z1with an objective water lens at 40× magnification. Images were processed using Fiji ImageJ software^43^. Images of mutant and wild-type loops were examined and the cell numbers were quantified.

### Reverse transcription PCR (RT PCR) and quantitative PCR (qPCR) analyses

Each fungal strain was cultured on 9 cm LNM Petri plates without supplement or with 10 mM NH_4_Cl and covered with cellophane in groups of 10 plates at 25°C for 7 days. Fungal hyphae were collected carefully and lyophilized. Total RNA was extracted using a Trizol-based protocol as described^44^ and purified using a Direct-zol RNA MiniPrep Kit (Zymo Research) according to the manufacturer’s instructions.

For cDNA synthesis, 2 μg pure RNA per 20μl reaction was reverse-transcribed using SuperScript IV Reverse Transcriptase (Invitrogen by Thermo Fisher Scientific) according to the manufacturer’s instructions, using oligo-dT-primed reverse transcription. Real-time qPCR was performed using a model QuantStudio 12K Flex Real-Time PCR System (Applied Biosystems by Thermo Fisher Scientific). Primers used for qPCR are listed in Supplementary Table 2.

Briefly, each 20μL reaction mixture contained 5ng cDNA, 100 nM each primer, and 10 μL Fast SYBR Green Master Mix (Applied Biosystems by Thermo Fisher Scientific). The reverse transcription process included an initial cycle at 95°C for 20 seconds, followed by 40 cycles of 95°C for 1 second and 60°C for 20 seconds. *GPD1* was used as an endogenous control for qPCR. Data from three replicates were used to estimate the mean ΔCT and standard deviation. The relative fold-change of each gene is shown according to the 2^−ΔCT^ method.

### Plasmid construction for heterologous gene expression

To express the *MEP* genes of *A. oligospora* in *S. cerevisiae*, we generated the vector pRS42N_OE using the *PGK1* promoter and terminator from *S. cerevisiae* in the multiple cloning sites of pRS42N plasmid containing restriction cutting sites between the promoter and terminator (Supplementary figure 6A). The cDNAs of the each *AoMEP* genes was amplified from total cDNA of *A. oligospora* after 4-hour nematode induction using Phusion High-Fidelity DNA Polymerase (Thermo Scientific). The pRS42N_OE vector was digested by HindIII and BamHI restriction enzymes (Thermo Fischer Scientific) and purified with a Gel Extraction Kit (QIAquick, QIAGEN). An In-Fusion HD Cloning Kit (Takara Bio) was used to fuse the vector and the cDNA fragment (Supplementary figure 6B). Plasmids exhibiting correct size after digestion were sent for Sanger sequencing (GENOMICS, https://www.genomics.com.tw/) to check cloning accuracy.

### Yeast transformation by electroporation

All yeast transformations were performed by using electroporation as described before with some modifications^45^. Yeast strains used for heterologous gene expression are listed in Supplementary Table 3. To generate yeast competent cells, yeasts were cultured in 10 mL YPD at 28°C overnight with shaking. The yeast cells were collected by centrifugation at 1500 rcf for 3 minutes and then mixed with 8 mL ddH_2_O, 1 mL 10× TE buffer, 1 mL 1 M lithium acetate and 250 μl 1 M DTT. The cells were then incubated at 30°C for 1 hour, before adding 30 mL cold ddH_2_O and then being spun down. The resulting pellet was washed with 5 mL cold 1 M sorbitol and resuspended in 0.5 mL cold 1 M sorbitol.

For each reaction, 200 ng of plasmid was mixed with 100 μl of yeast competent cells and transformed by electroporation. After electroporation, the cells were mixed with 1 mL sorbitol and transferred to a 1.5-mL Eppendorf for 3-hour recovery at room temperature. Finally, the cells were plated on YPD containing 75 μg/mL clonNAT and incubated at 28°C for 2 to 3 days.

### Functional analysis of AoMep1, AoMep2, and AoMep3 in *S. cerevisiae*

To assess whether AoMep transporters could complement the pseudohyphal growth defect of the *S. cerevisiae mep2* mutant (MLY108a/*⍺*) under low ammonium conditions, plasmids containing *AoMEP* cDNA were transformed into the *Scmep2* mutant. As controls, the empty vector pRS42N_OE was transformed into both the wild-type strain (MLY97a/*⍺*) and the *Scmep2* mutant. The strains were initially grown on YPD plates at 28°C for 2 days, after which single colonies were streaked onto SLADG agar medium. After incubation at 30°C for 6 days, colony images were captured using a ZEISS Stemi 305 Stereo Microscope at 80× magnification.

To determine if AoMep transporters could rescue the growth defect of the *S. cerevisiae mep1 mep2 mep*3 triple mutant (MLY131a/*⍺*) under low ammonium conditions, plasmids encoding *AoMEP* genes were introduced into the triple mutant. The pRS42N_OE empty vector was also transformed into both the wild-type strain (MLY97a/*⍺*) and the triple mutant as controls. Single colonies grown on YPD were transferred to YPD liquid medium for overnight culture at 28°C. The cell density was adjusted to a final OD600 of 0.2 by diluting in 120 μl of SLADG medium containing 25 μM ammonium sulfate in 96-well plates. The plates were incubated at 28°C for 2 days and analyzed using a plate reader (Infinite 200 PRO, TECAN).

## Data Availability Statement

The authors affirm that all data necessary for confirming the conclusions of the article are present within the article, figures, and tables. Strains and plasmids are available upon request.

## Acknowledgments

We thank Dr. Joseph Heitman for sharing *S. cerevisiae* strains. This study is supported by Academia Sinica Investigator Award AS-IA-111-L02 (YPH) and Max Planck Society (YPH).

## Supplementary tables and figures

**Supplementary table 1.**
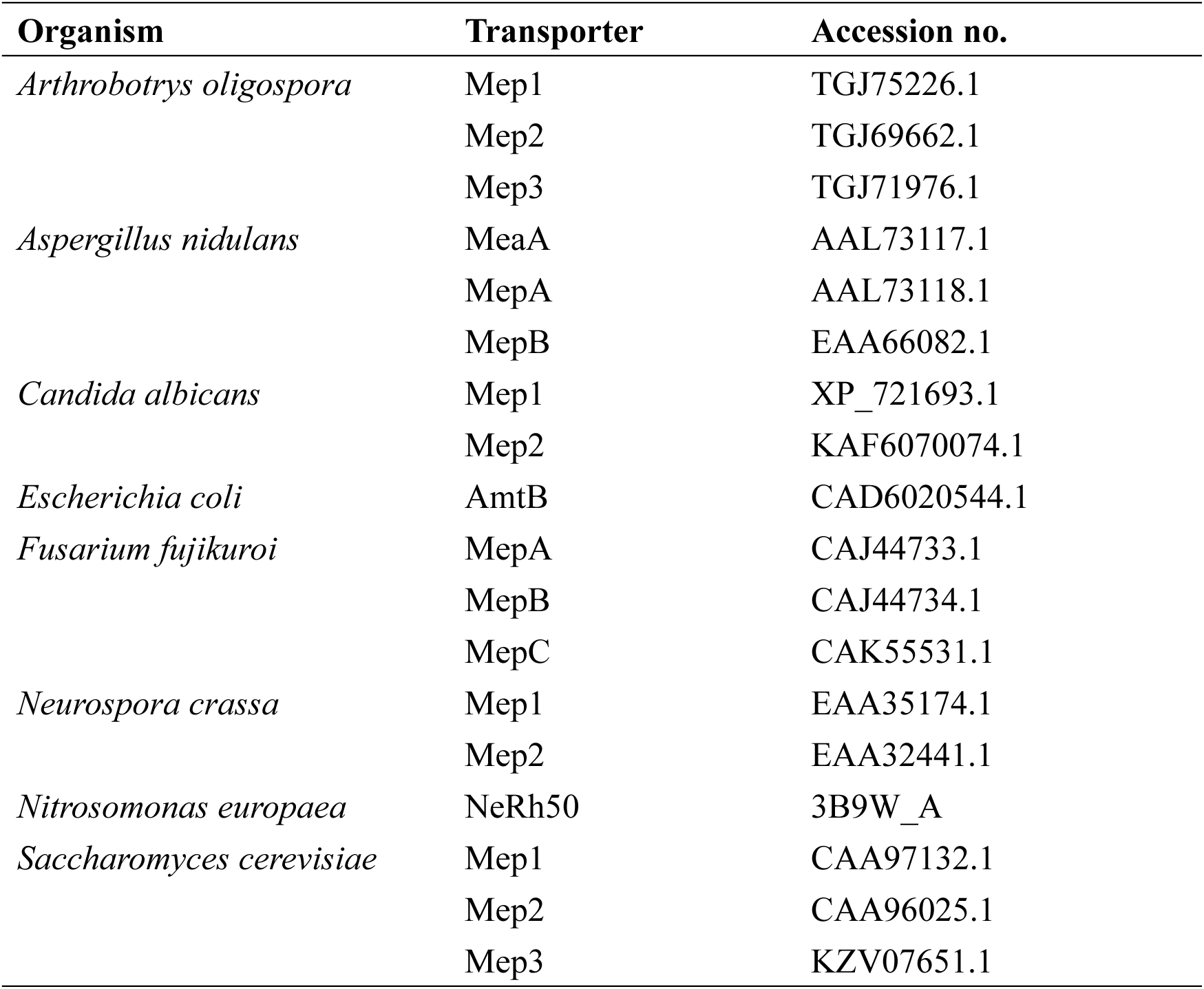
Ammonium transporters of different organisms used for phylogenetic analysis.

| Organism | Transporter | Accession no. |
| --- | --- | --- |
| *Arthrobotrys oligospora* | Mep1 | TGJ75226.1 |
|  | Mep2 | TGJ69662.1 |
|  | Mep3 | TGJ71976.1 |
| *Aspergillus nidulans* | MeaA | AAL73117.1 |
|  | MepA | AAL73118.1 |
|  | MepB | EAA66082.1 |
| *Candida albicans* | Mep1 | XP_721693.1 |
|  | Mep2 | KAF6070074.1 |
| *Escherichia coli* | AmtB | CAD6020544.1 |
| *Fusarium fujikuroi* | MepA | CAJ44733.1 |
|  | MepB | CAJ44734.1 |
|  | MepC | CAK55531.1 |
| *Neurospora crassa* | Mep1 | EAA35174.1 |
|  | Mep2 | EAA32441.1 |
| *Nitrosomonas europaea* | NeRh50 | 3B9W_A |
| *Saccharomyces cerevisiae* | Mep1 | CAA97132.1 |
|  | Mep2 | CAA96025.1 |
|  | Mep3 | KZV07651.1 |

**Supplementary table 2.** Oligonucleotides used in this study.

*Excel file

**Supplementary table 3.**
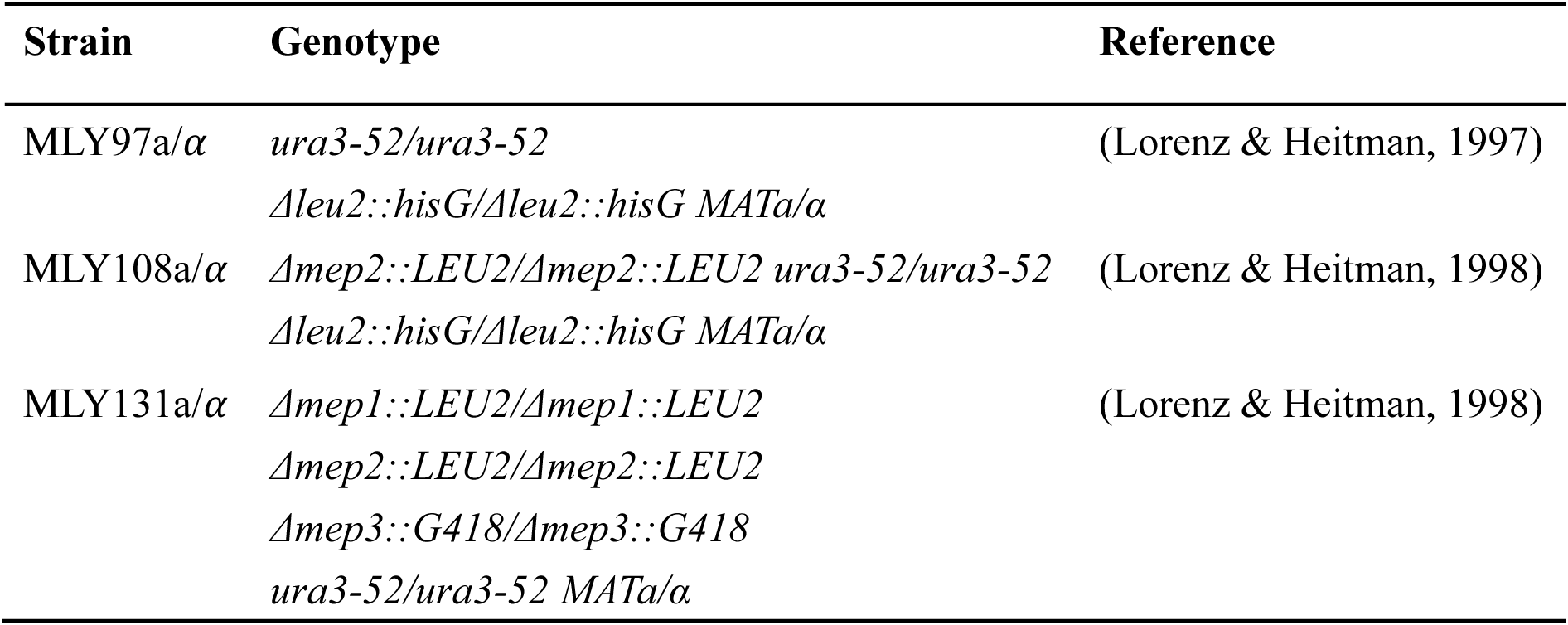
*Saccharomyces cerevisiae* strains used for heterologous gene expression.

| Strain | Genotype | Reference |
| --- | --- | --- |
| MLY97a/$\alpha$ | *ura3-52/ura3-52*<br>*Δleu2::hisG/Δleu2::hisG MATa/$\alpha$* | (Lorenz & Heitman, 1997) |
| MLY108a/$\alpha$ | *Δmep2::LEU2/Δmep2::LEU2* *ura3-52/ura3-52*<br>*Δleu2::hisG/Δleu2::hisG MATa/$\alpha$* | (Lorenz & Heitman, 1998) |
| MLY131a/$\alpha$ | *Δmep1::LEU2/Δmep1::LEU2*<br>*Δmep2::LEU2/Δmep2::LEU2*<br>*Δmep3::G418/Δmep3::G418*<br>*ura3-52/ura3-52 MATa/$\alpha$* | (Lorenz & Heitman, 1998) |

**Supplementary figure 1.**
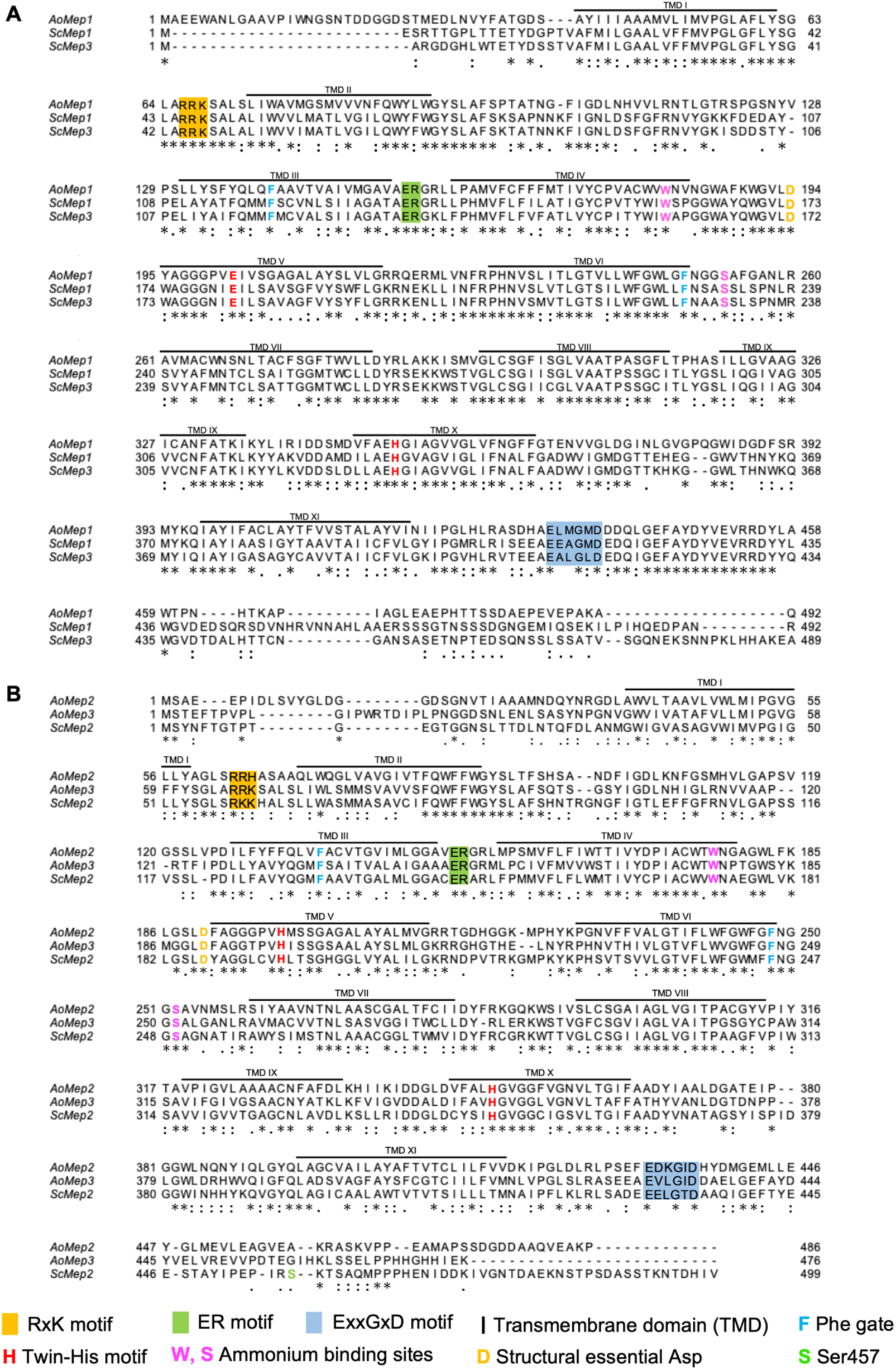
Protein sequence alignment of the ammonium transporters between *Arthrobotrys oligospora* and *Saccharomyces cerevisiae*. The protein sequences were aligned using online MAFFT (multiple alignment using fast Fourier transform) program ver.7^31^ (https://mafft.cbrc.jp/alignment/server/). (A) Alignment of AoMep1, ScMep1, and ScMep3. (B) Alignment of AoMep2, AoMep3, and ScMep2. Asterisks (*) indicate identical amino acids, colons (:) indicate conserved amino acids and single dots (.) indicate semi-conserved amino acids. Predicted transmembrane domains (TMDs) are shown above the alignment. Conserved motifs are shown in colored boxes, including the RxK motif in intercellular loop1 (ICL1) (yellow), the ER motif in ICL2 (green), and the ExxGxD motif in the C terminus (blue). Key residues are shown as colored letters, including the Phenylalanine gate (blue), the Tryptophan binding site (magenta), the twin-Histidine motif (red), and the Npr1 phosphorylation target Serine457 in the ScMep2 C terminus (green).

**Supplementary figure 2.**
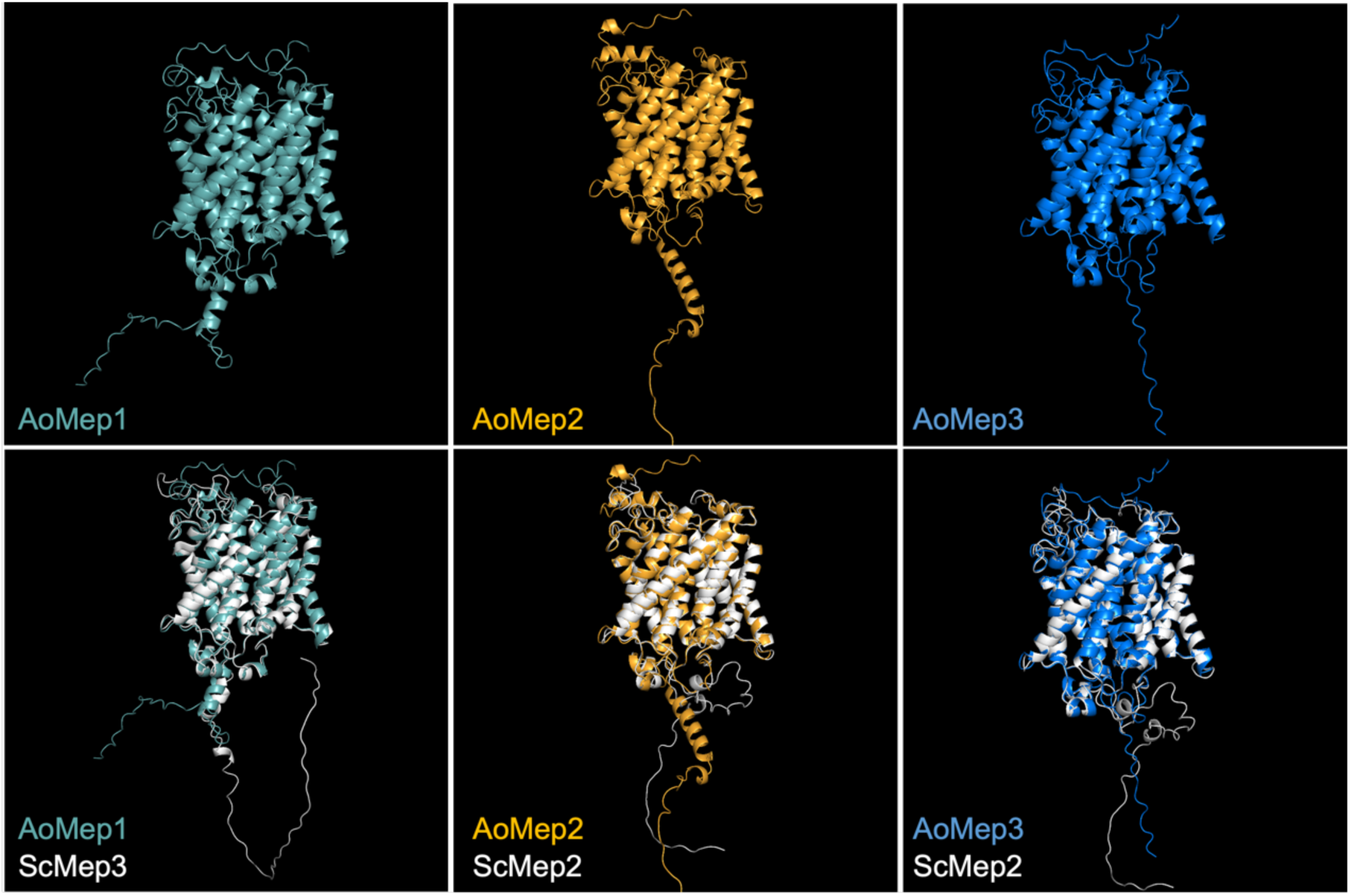
Predicted protein structures of AoMep transporters and the structure alignment to ScMep transporters. Protein structures were predicted using AlphaFold^35^, structural alignments were performed using Foldseek^36^, and molecular visualizations were generated using PyMOL^37^. The upper panel shows the structures of individual AoMep transporters, while the lower panel displays their structural alignment with the *S. cerevisiae* Mep transporters (shown in white).

**Supplementary figure 3.**
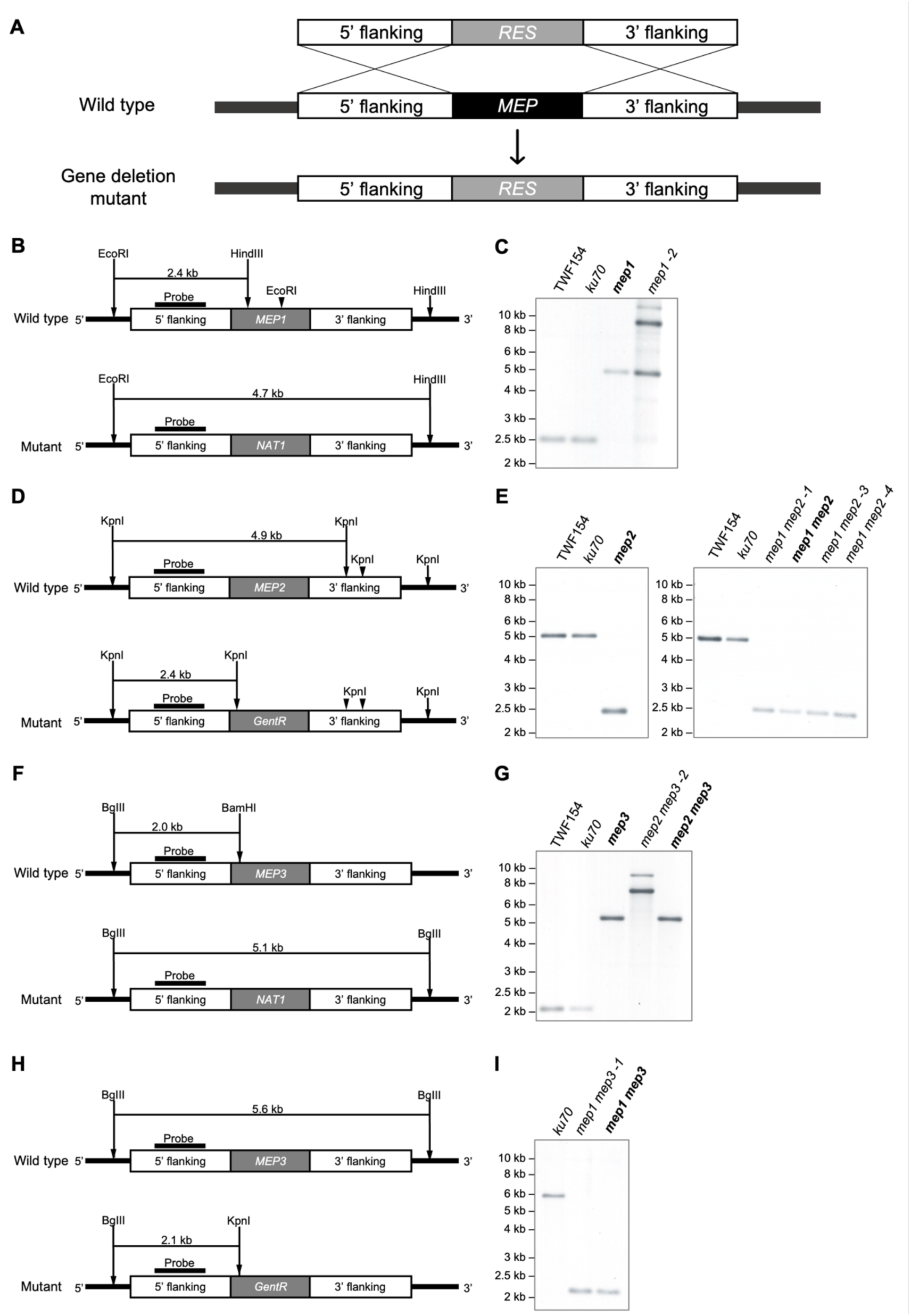
Schematic of *A. oligospora mep* gene deletion by homologous recombination and Southern blot confirmation. (A) Endogenous *MEP* genes were deleted by homologous recombination using constructs containing homologous 5’ and 3’ flanking regions fused with either *NAT1* or *GentR* antibiotic resistance genes (“*RES*”). (B, D, F, and H) Schematics showing the restriction enzyme cutting sites and probes for both the wild-type and mutant constructs. Restriction enzyme cleavage sites, fragment sizes, and resistance genes are marked in each figure. The probes were chosen to hybridize to the 5’ flanking regions of the cassettes. (C, E, G, and I) Results of Southern blotting according to the schematics on the left. Wild-type (TWF154) and gene deletion background (*ku70*) strains were used as controls. Mutant strains used in further experiments are indicated in Southern blots using bold text.

**Supplementary figure 4.**
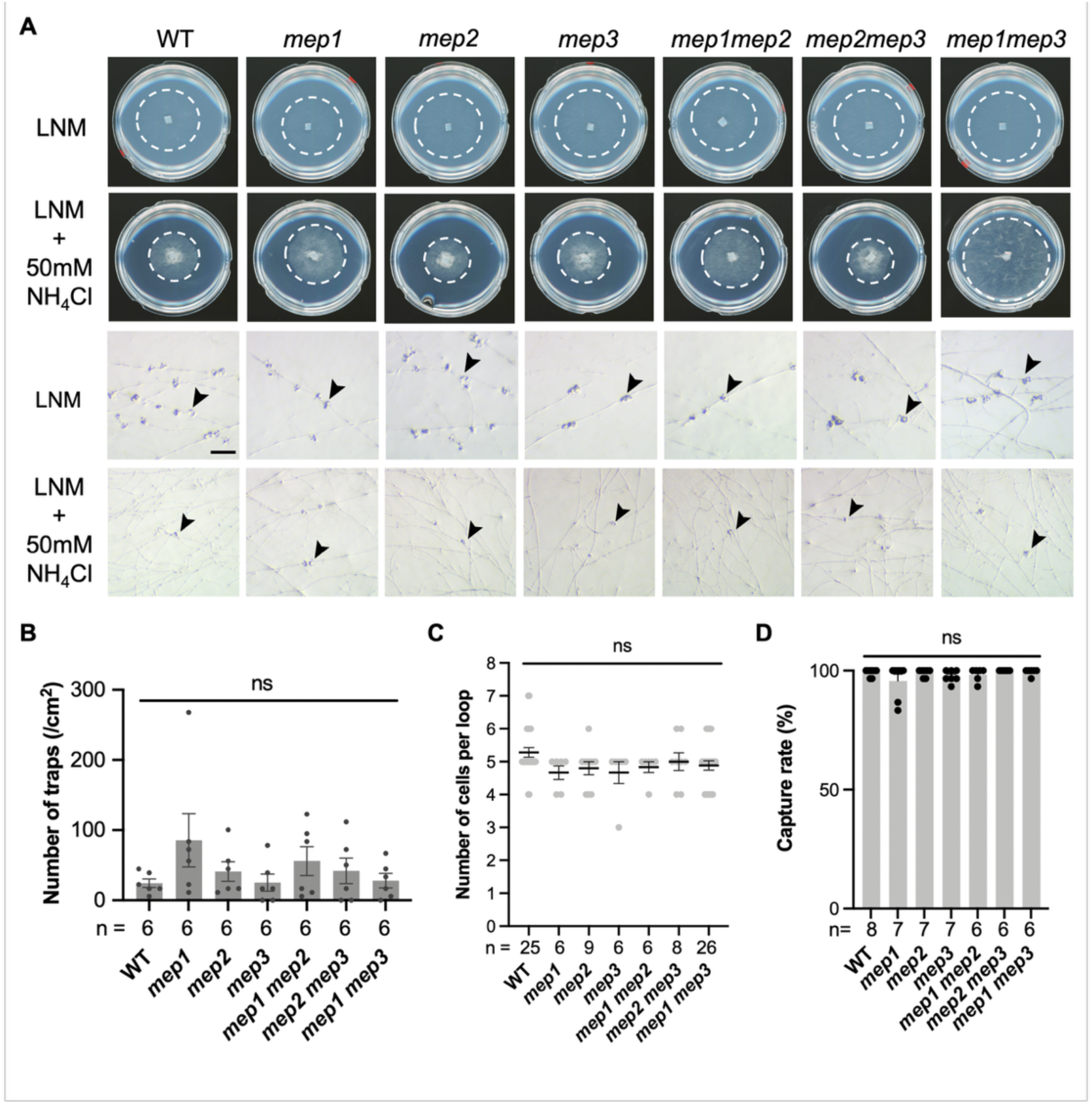
Phenotypic assay of *mep* mutant strains. (A) Hyphal growth and representative brightfield images of the traps induced by nematodes in the WT and *mep* mutants on LNM supplemented with 50 mM ammonium chloride. Dashed lines indicate the colony edges. Arrowheads indicate representative traps. (B) Quantification of trap formation in *(A)*. Data are presented as mean ± SEM. Comparisons within groups were analyzed by two-way ANOVA followed by Dunnett’s multiple comparisons test. (C) Quantification data for cell numbers per trap loop. There is no significant difference in cell number between mutants and the wild type as detected by ordinary one-way ANOVA (*P*>0.05). (D) Capture rate assay. No significant difference (ns) was seen as analyzed by ordinary one-way ANOVA, showing that the traps of *mep* mutants are still functional. WT, wild type.

**Supplementary figure 5.**
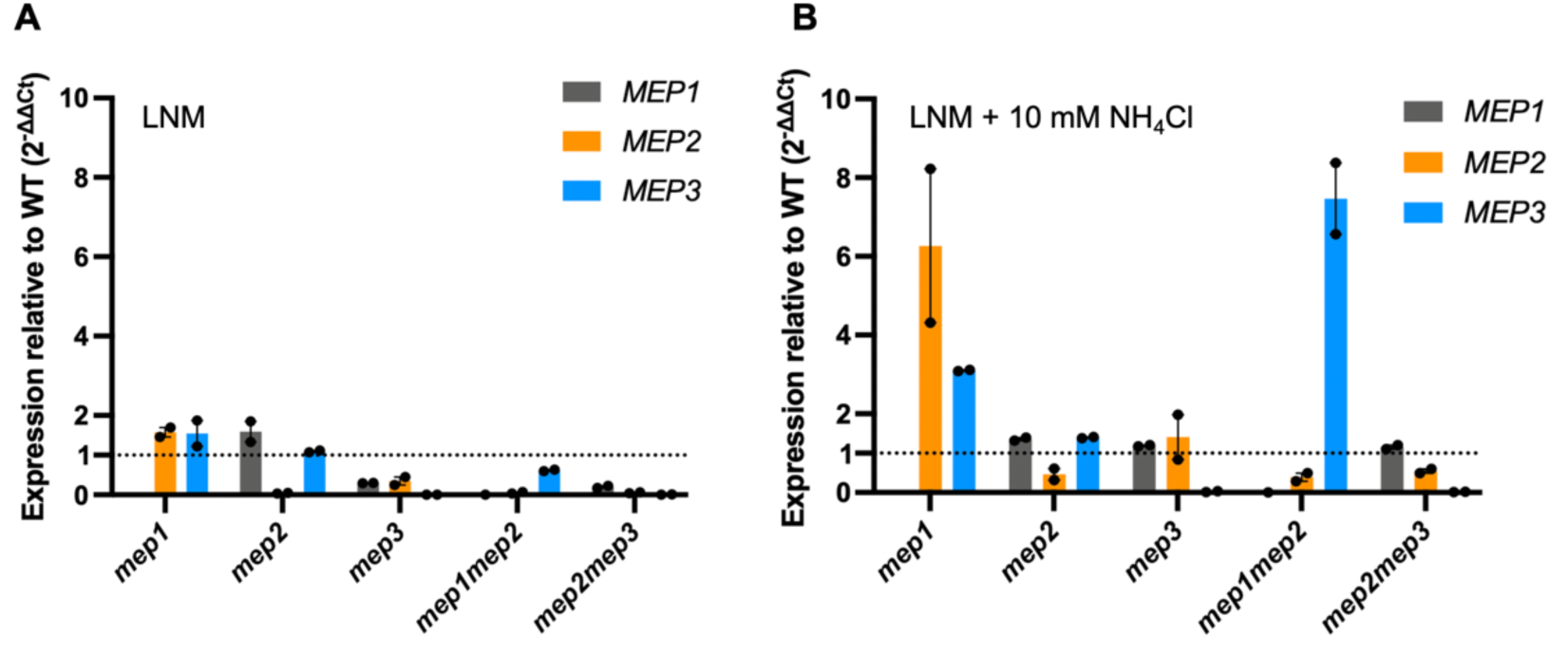
Relative *MEP* gene expression in *mep* mutants in response to the presence of ammonium using quantitative PCR. *MEP* gene expression was quantified in *mep* single and double mutants, normalized to *GDP1*, and expressed using 2^−ΔΔCt46^, meaning that at 1 on the Y-axis relative expression of the measured gene should be unchanged as compared to the wild type in the same conditions. Fungi were grown on low-nutrient medium (LNM) (A) or on LNM supplemented with 10 mM NH_4_Cl (B).

**Supplementary figure 6.**
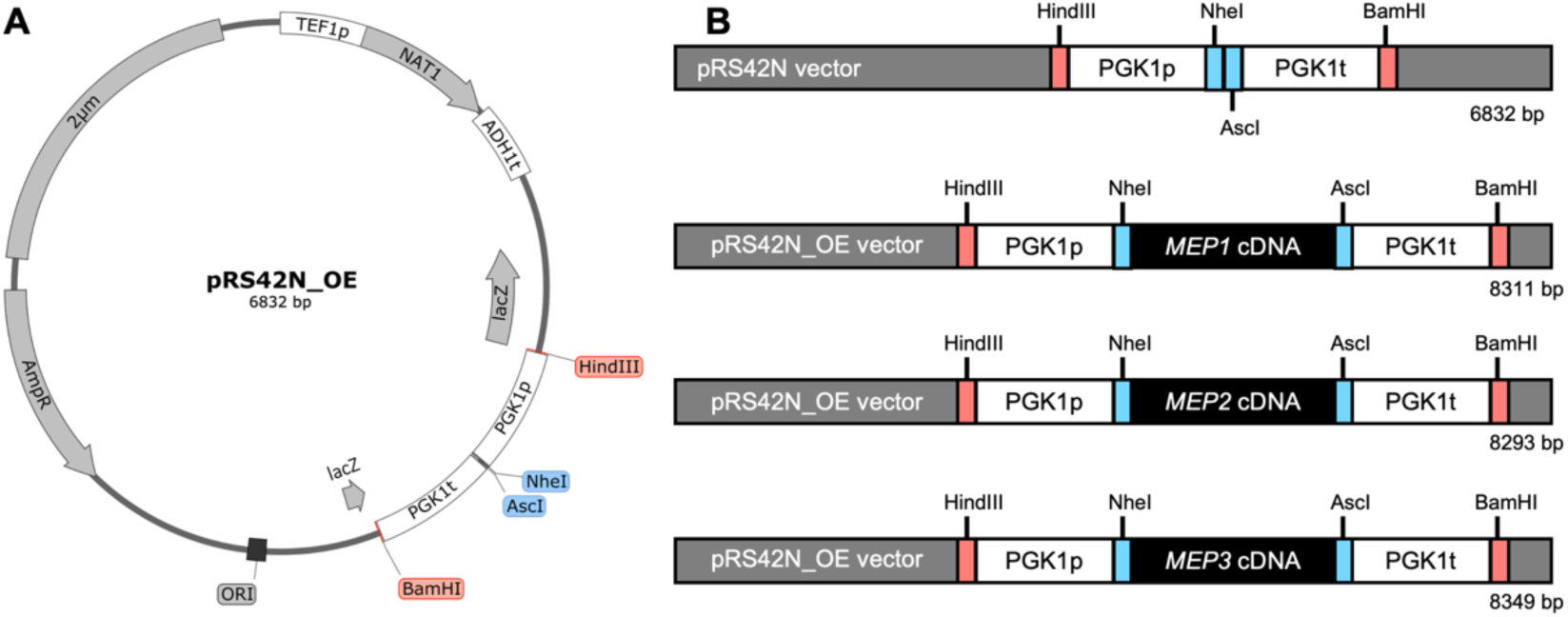
Plasmid map of pRS42N_OE and heterologous gene expression maps of *AoMEP* genes. (A) The pRS42N_OE plasmid was generated based on a pRS42N backbone with the *S. cerevisiae PGK1* promoter and *PGK1* terminator inserted at the multiple cloning site using the BamHI and HindIII restriction sites. cDNAs were inserted between the *PGK1* promoter and terminator using the restriction sites AscI and NheI. (B) Schematic diagrams of the pRS42N_OE plasmid and *AoMEP* heterologous gene expression plasmids. The cDNAs of *AoMEP* were amplified from total cDNA of 4-hour nematode-induced *A. oligospora* (TWF154), and integrated between the NheI and AscI restriction sites. Total plasmid lengths are shown at the bottom right of each map.

**Supplementary figure 7.**
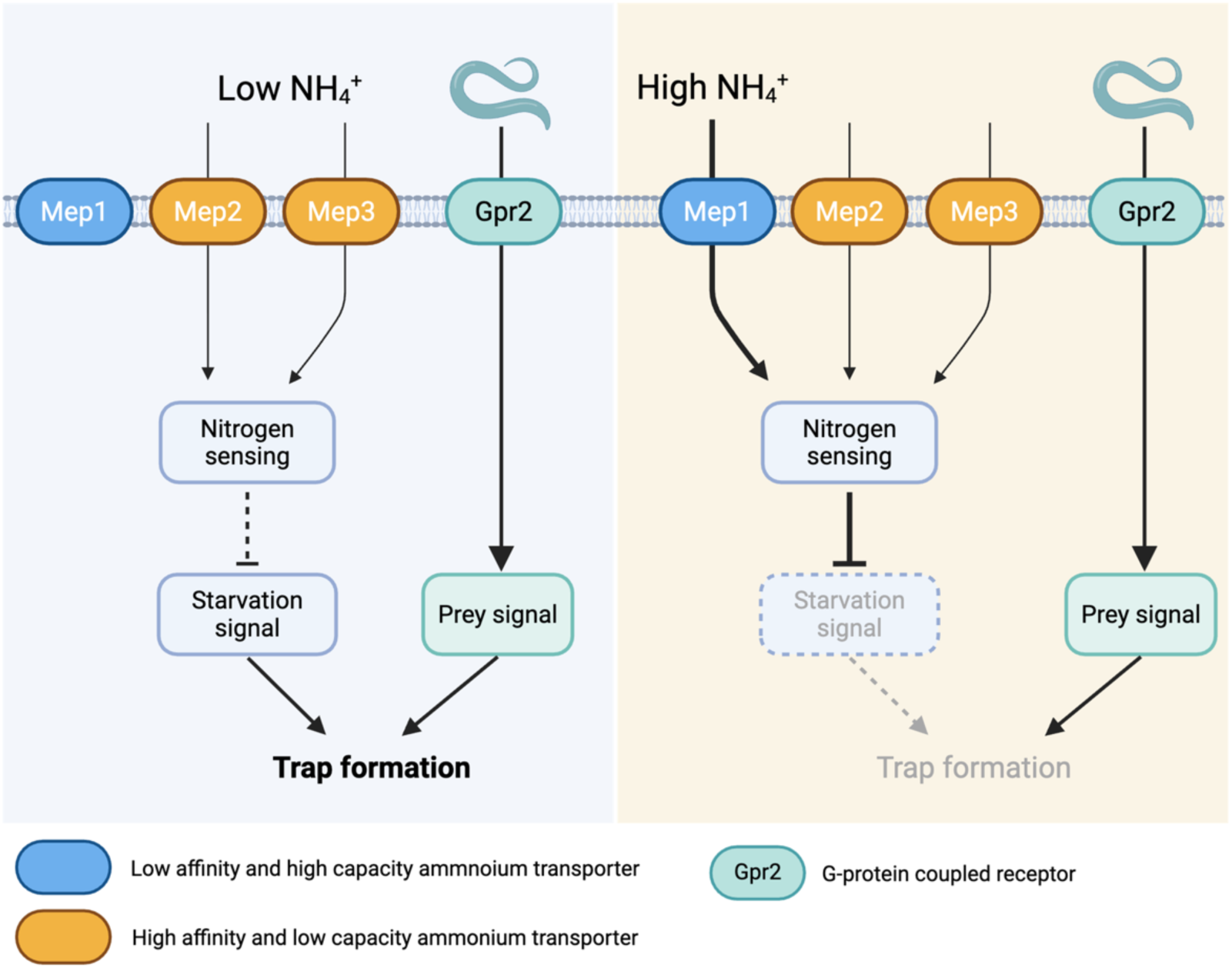
Model of how Mep transporters operate in *A. oligospora*. Under low ammonium condition, Mep2 and Mep3 play roles in facilitating ammonium transport into fungal cells. *A. oligospora* perceives nutrient deprivation to produce a signal that, when coupled with prey signals generated by the GPCR protein Gpr2, initiates trap morphogenesis. In environments with high ammonium levels, Mep1-mediated transport leads to elevated intracellular ammonium concentrations, signaling a non-starvation state that suppresses trap formation despite the detection of prey signals.

